# Modeling *Edar* expression reveals the hidden dynamics of tool signaling center patterning

**DOI:** 10.1101/453258

**Authors:** Alexa Sadier, Monika Twarogowska, Klara Steklikova, Luke Hayden, Anne Lambert, Pascal Schneider, Vincent Laudet, Maria Hovorakova, Vincent Calvez, Sophie Pantalacci

## Abstract

The generation of patterns during development is generally viewed as a direct process. In the mouse jaw, however, the sequential patterning of molars initiates with abortive tooth signaling centers called MS and R2, thought to be vestiges of the lost rodent premolars. Moreover, the mature signaling center of the first molar (M1) is formed from the fusion of two signaling centers (R2 and early M1). Here, we report that *Edar* expression reveals the hidden dynamics of signalling centers patterning. First, *Edar* expression evidenced a hidden two-step patterning process that we modelled with a single activator-inhibitor pair: the epithelium is initially broadly activated, then activation becomes restricted in space to give rise to the signalling centers. Second, *Edar* expression unveils successive phases of pattern making and pattern erasing events, a phenomenon that we called a developmental palimpsest. MS is erased by a broad activation for the benefit of R2, which itself is erased before it recovers when the first molar signaling center forms. In the lower but not the upper jaw, the two neighboring signaling centers then fuse into a single elongated center. Our model recapitulated the erasure of the R2 signaling center by the wave of activation that precedes the formation of M1 signaling center, and predicted the surprising rescue of R2 in the context of an *Edar* mutant with reduced activation. It suggested that R2 was not intrinsically defective, but actively outcompeted by M1 formation. We confirmed this by cultivating R2 separately from the posterior tissue and showing it could then generate a tooth. Finally, by introducing chemotaxis as a secondary process of tooth germ maturation, we recapitulated the fusion of R2 and M1 in the lower jaw only, and the loss of fusion when Edar function is impaired in organ cultures. In conclusion, we have uncovered a highly indirect and dynamic nature of pattern formation in the molar field that could nevertheless be simulated with simple mathematical models. Our study argues for viewing embryonic patterns as dynamical objects rather than as fixed endpoints, where dynamics is essential to the outcome of the patterning process.

## Introduction

The emergence of ordered patterns in multicellular organisms has been a major field of research in developmental biology, revealing a diversity of pattern formation mechanisms. While some patterns appear simultaneously (e.g. drosophila segments, mouse hair), other appear sequentially (e.g. feathers on chicken’s back), most often as the structure grows distally (e.g. short-germ insects segments, somites, limbs proximo-distal elements, palatal rugae). Several types of patterning mechanisms have been proposed, some relying on a prepattern (e.g. “positional information” model: a gradient of a signaling molecule is turned into a more complex pattern by interpreting the varying concentration at each position in space, (1,2)) and other on spontaneous pattern formation (e.g. reaction-diffusion (Turing) mechanisms, chemotaxis, see below and (3–5)). In these studies, more or less importance has been given to the temporal dynamics of pattern formation depending on the mechanism. Sequential formation requires the consideration of temporal aspects that can be neglected when the pattern forms at a glance (6,7). Spontaneous pattern formation inherently emphasizes the dynamics of the system. Instead, positional information has been mostly associated with static representations (e.g. the French flag model, see (3)). In all cases however, patterning is viewed as a conceptually simple temporal process: from a prepattern or a spatial heterogeneity emerges the final pattern. It is however questionable whether biological systems, which result from an historical, contingent process, proceed in such a direct manner, or if transient patterns can be constructed and deconstructed during embryogenesis until the final pattern is formed. Recently, the textbook example of simultaneous pattern formation, i.e. the formation of *Drosophila* gap gene expression pattern, was closely reexamined, and it was found that the gene expression pattern was changing with time, as maternal inputs decay, with important consequences for the final pattern (8). To our knowledge, other examples are lacking. Here, we studied this question in a model of sequential patterning: the mouse molar row.

The search for the general mechanisms generating patterns in biology has been greatly influenced by the theoretical work of the mathematician Alan Turing (4,5,9). The generalization of this work has led to many classes of reaction-diffusion (RD) mechanisms, where two (or more) molecules characterized by different spatial range of action and a given topology of interaction can self-organize a stable pattern, but also exhibit behaviors such as oscillations or propagating waves (4). The most iconic example is the case where a short-range activator that self-amplifies and activates its own long-range inhibitor, can create spots, stripes or labyrinths. Recently it has been shown that many biological systems exhibit features of RD mechanisms (e.g. patterning of epithelial appendages such as hair (10,11), the patterning of features such as the rugae of the palate (12), digits (13) and somites (14)). This should not be taken too strictly however. Geirer and Meinhardt pointed out that any process involving local self-enhancement and lateral inhibition has the potential to drive spontaneous pattern formation (15). For example, color pattern formation in zebrafish can be explained by RD models, but at least partly involves cell interactions rather than the diffusion of biomolecules (16,17). Pattern formation can also arise from purely chemostaxis mediated self-organization. When cell movement is driven by concentration gradients of chemostatic cues, positive feedbacks between cell density and chemo-attractant production are known to enhance local concentration of cells and may result in self-sustained aggregation. (18–20). Chemotaxis plays a prominent role in feather formation (21), and this is likely also the case in most other epithelial appendages (e.g. hair (Glover et al., 2017)).

Mouse molars are a good example of repeated structures that form through sequential pattern formation. Mice have only three molars per quadrant, separated from incisors by a diastema, the other mammalian teeth (i.e. canine and premolars) having been lost in the evolution of mouse lineage (22). Molars develop sequentially from the most anterior molar (first molar, M1) to the most posterior (third molar, M3). They develop from a unique cylinder-shaped invagination of the oral epithelium, the so-called dental lamina, (23–25) where tooth-specific signaling centers, called Primary Enamel Knots (PEK) are patterned. These signaling centers then drive the formation of individual teeth by promoting “cap” formation; the capping of the underlying condensed mesenchyme by the epithelium. Indirect evidence that activation-inhibition mechanisms determine sequential formation of these signaling centers comes from the similarity of tooth formation with other epithelial appendages (26), namely hair and palatal rugae, whose patterning in clearly ruled by Turing-type mechanisms ((11,12)). The most direct evidence is a study by Kavanagh and colleagues (27), showing that when tissue that will form M2 is separated from M1, M2 forms earlier and becomes larger. PEK formation in the epithelium requires signaling from both the epithelium and the mesenchyme (28), including a mechanical signal induced by mesenchyme condensation (29).

The sequential patterning of mouse molars in the lower jaw (Figure 1A) involves two transient signaling centers (30), that fail to drive proper cap transition, yet form morphologically distinguishable buds (30,31). These buds-might be the vestige of lost premolars (30,32). Monitoring these signaling centers *via Shh* expression has shown that the signaling center called MS initiates sequential patterning and then disappears (30). Subsequently, the R2 signaling center forms. As it vanishes, the M1 early signaling forms posteriorly (Prochazka et al., 2010). Soon after, the former R2 and M1-early signaling centers are encompassed in a giant -*Shh*-expressing signaling center (Prochazka et al., 2010; Lochovska et al., 2015). Here, we will refer to it as the mature M1 signaling center. A similar situation with two abortive buds (called R1 and R2) has been noticed in the upper jaw (25). Their signaling centers have not yet been characterized, although they are morphologically more apparent than in the lower jaw.

**Figure 1.**
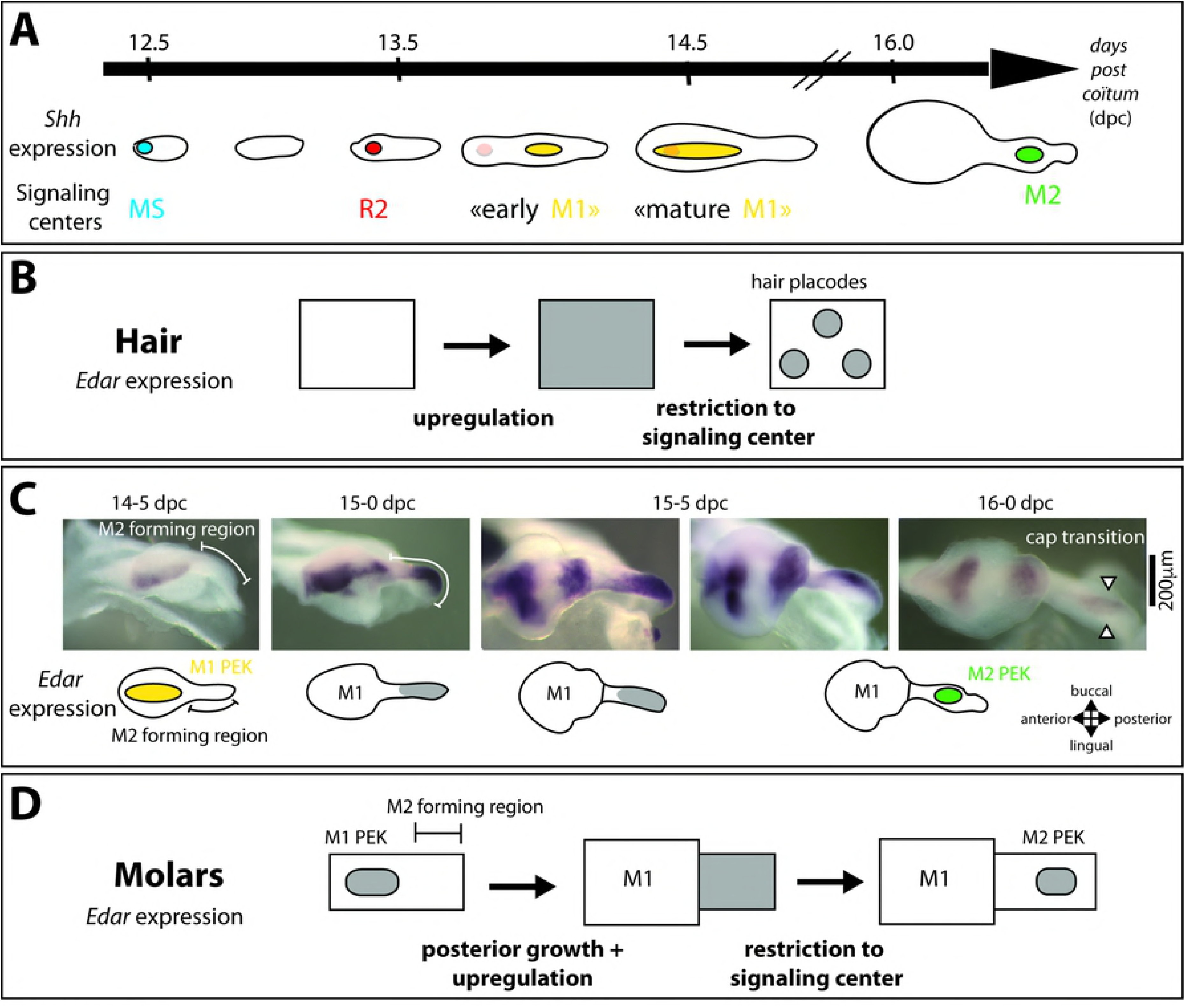
***Edar* expression is dynamically regulated during hair and tooth patterning** A- Scheme summarizing the sequential patterning of signaling centers in the dental lamina as revealed with *Shh* expression. B- Scheme showing *Edar* regulation in epidermis during primary hair patterning. Concomitantly with hair placodes patterning, *Edar* becomes upregulated in the hair placode while being downregulated in its neighborhood, highlighting the outcome of the activation-inhibition mechanisms that patterned the placode and avoid further formation of placodes in the vicinity. C- *In situ* hybridization showing *Edar* expression in the isolated dental epithelium from 14.5 to 16.0 days, a period corresponding to growth of the tail region (“M2 forming region“) and patterning of M2 signaling center in this newly grown region (at 16.0). At 14.5, *Edar* is restricted to the primary enamel knot of the first molar (yellow on the scheme). At 15.0, *Edar* expression is seen in the posterior-most part of the tail, the M2-forming region (Gray on the scheme). Between 15.5 and 16.0, it is restricted to the primary enamel knot of the second molar (M2, green on the scheme). On the scheme, the continuing expression of *Edar* in the maturating first molar has been omitted for simplicity. D- Scheme summarizing the expression pattern in C, to be compared with B. The situation in hair is similar to the situation in the anterior part of the dental epithelium.

Interestingly, mutations in genes affecting various developmental pathways (FGF, Shh, Wnt, BMP and Eda pathways) lead to a supernumerary tooth in front of M1, thus resembling a premolar (34). Where it was specifically studied, the results were consistent with R2 signaling center being enabled to form a tooth (35–39,33). The picture is thus fairly complex, especially since we lack direct evidence for the dynamics of activation-inhibition mechanisms that pattern signaling centers in the dental lamina and promote tooth formation.

The Eda pathway has the potential to shed light on these mechanisms. This pathway shows a consistent role in activation-inhibition mechanisms throughout several epithelial appendages (hair, feather, teeth; (40,41). This role has been more specifically evidenced for mouse guard hair pattern formation. The receptor of the pathway, Edar, is first broadly expressed in the epidermis. Concomitantly with hair signaling center patterning, it becomes upregulated in the placodal signaling center, and downregulated in the neighborhood (Figure 1B). In the absence of Edar signaling, no signaling center forms (11,42,43), while increasing signaling results in more numerous and densely packed placodes (11). Current models posit that the Eda pathway is activated by Wnt, activin BA and BMP4 pathways (43,44) (11,44), but also feeds back on these pathways (and others) through the transcriptional activation of their diffusing ligands and inhibitors (*e.g*. WNT10a/b, DKK4, CCN2/CTGF, Follistatin and FGF20 (40,41)). More recently, it appeared that Eda signaling also promotes placodal fate by stimulating the centripetal migration of cells in the epithelium (45,46). In teeth, the Eda pathway is dispensable for primary signaling center (PEK) formation, but required for its correct sizing (46–48). Similarly, it is necessary for correct patterning of the secondary signaling centers controlling cusp morphogenesis (38,47,49). *Eda* and *Edar* mutants have reduced tooth size and cusp number, but intriguingly, sometimes have a small supernumerary tooth (50–52). In gain-of function mutations, an anterior supernumerary tooth is also found, and teeth are larger with more cusps (49,53–55).

In this paper, we aimed at clarifying the temporal dynamics of signaling center formation in the dental lamina. We studied the temporal dynamics of *Edar* gene expression, the receptor of the Eda pathway, during molar pattern formation and showed it recalls the dynamics observed during hair patterning. Based on these data, we built a reaction-diffusion type model of molar patterning that enables sequential signaling center formation and helps reveal the exquisitely complex temporal interactions leading to the construction and deconstruction of patterns in the developing molar row. Our model explains a counter-intuitive result, the rescue of the abortive R2 bud in the context of reduced activation/increased inhibition of the Edar loss of function mutation. Finally, we show that Edar is necessary for the formation of a fused R2-M1 signaling center in the lower jaw only, possibly through a chemotactic effect. We thus showed that patterning is not direct, although it follows simple mathematical rules.

## Results

### *Edar* regulation highlights activation in the growing dental epithelium

To get insights into molar row patterning, we examined the regulation of the *Edar* gene (Figure 1B). Because the early period of molar row patterning is complicated by the presence of vestigial signaling centers, we first focused on the patterning of the second molar. Since *Edar* is exclusively expressed in the epithelium, we performed *in situ* hybridization on mandibular epithelium that has been dissociated from the mesenchyme, thus providing a 3D view of *Edar* expression. At 14.5 dpc, *Edar* expression is restricted to the primary signaling center (PEK) of the first molar, and no expression is seen in the second molar field, looking like a “tail” (Figure 1C). At 15.0 dpc, the “tail” has elongated and *Edar* expression is upregulated in the posterior most part of it. By late 15.5 dpc, it starts restricting to the M2 primary enamel knot, just before M2 cap transition occurs (at 16.0 dpc). The restriction was concomitant with *Shh* expression starting in M2 PEK (data not shown and (30)).

This dynamic of an initial broad upregulation of *Edar* followed by its restriction to a signaling center is reminiscent of what happens during hair patterning (compare Figure 1B and 1D). It suggests that the decision to form a tooth signaling center in the growing molar field proceeds in two phases. First, the whole dental epithelium is activated. This activation results in broad *Edar* expression, so far the only gene to show such an expression pattern marking the epithelium competent to form a tooth. Second, activation gets restricted spatially and gives rise to a signaling center. This results in the focused expression of Edar, and many other genes known as primary enamel knot genes (*e.g. Shh)*. In this view, *Edar* expression is a read-out of activation levels in the molar field: where it is high enough, *Edar* is expressed. To further formalize these ideas, we built a mathematical model of activation in the dental epithelium, as followed by *Edar* expression pattern.

### Activation dynamics can be modeled by a transition from a bistable system to a Turing system in a growing domain

Activation (monitored by Edar expression) can switch between different states: from no activation (that is no *Edar* expression) to broad activation (broad *Edar* expression) and from broad activation to spatially-restricted activation (focused *Edar* expression) suggestive of Turing mechanisms in the dental lamina. From a mathematical point of view, these complex behaviors can be modeled as two regimes of the same reaction-diffusion system (Figure 2A). The first regime, named throughout this paper *the bistable regime*, describes solutions connecting two constant, stable states (respectively, a high activation state and a low (no) activation state). This corresponds to the anterior part of the tissue with transient up-regulation of *Edar*. Second, the so-called Turing regime, differs from the previous one by the modification of one parameter (auto-inhibition strength). It is characterized by stable heterogeneous patterns which emerge from homogeneous patterns (*e.g*. spots). Thus the same system of equations describing the interaction of an activator and its inhibitor models waves of activation in the dental lamina as well as its restriction to the signaling center. A simple change in one reaction parameter could switch the system from the bistable region to the Turing regime. Because the restricted expression is only seen in the developmentally advanced parts (i.e. the anterior part) and follows broad activation, we impose that the activator has a positive feedback on tissue maturation, resulting in the switch to Turing regime. Biologically, this means that upon the broad wave of activation, new gene products have been produced that modify the activation-inhibition parameters.

**Figure 2.**
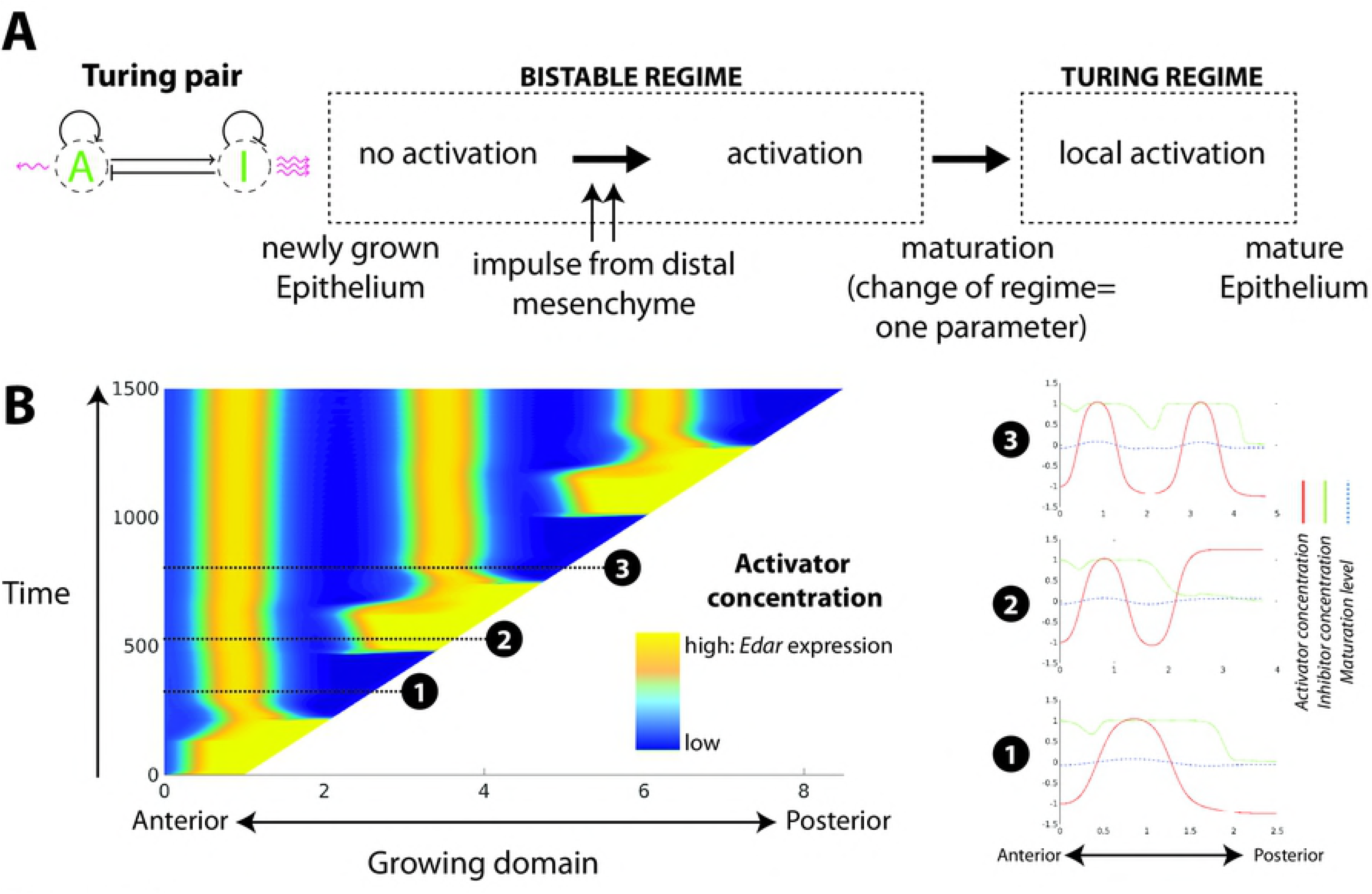
**a model for signaling center patterning based on maturation-dependent shift from a bistable to a Turing regime** A- Basic principles for a mathematical model of activation in the dental field, with *Edar* expression considered as a read-out of activator concentration. The dental epithelium first transits from no activation to generalized activation, as the tissue is primed by the anterior mesenchyme. Activation induce tissue maturation, moving the system to a Turing regime, which needs a single parameter change. Activation is localized to the signaling center. B- We modeled activation-inhibition mechanisms along the antero-posterior axis (1 dimension). Activator concentration is shown through space (x) and time (y), as the domain grows. Snapshots taken at three different timepoints are shown (red: activator concentration, blue: inhibitor concentration, green: domain maturation). The simulation shows the periodic behavior of the model: The dental epithelium grows in an inactivated state, due to inhibition coming from the Turing spot (snapshot 1, see also Figure 1C, 14.5 dpc). Newly grown parts of the dental epithelium get activated on a periodic basis (bright yellow anterior domains, snapshot 2, figure 1C 15.0–15.5 dpc), and upon maturation, produce a Turing peak (snapshot 3, Figure 1C 16.0 dpc). The movie corresponding to the snapshots is available as supplementary material 2.

Based on the above assumptions, we built a one-dimension model of RD mechanisms in an oriented growing domain. The model describes the time evolution of concentration of an activator and its inhibitor, which diffuse with different speeds and undergo kinetic reactions with explicit parameters regulating the transition between Turing and non-Turing regimes (Figure 2A). A detailed description and the parameters used in all simulations are found in the supplementary material (Supplementary Material 1). Below we summarize the main characteristics of the model, that exhibits periodic behavior, as shown in a representative simulation (Figure 2, see also the movie as Supplementary material 2).

The tissue grows from its anterior end, and the newly produced tissue matures exponentially in time (albeit at a slow rate). Maturation is stimulated in presence of the activator, with a certain time delay. In zones where maturation reaches a threshold value, the system parameters irreversibly switch from a bistable to a Turing regime (Figure 2B, snapshots 2 to 3).

Before this switch can occur, such simple system first needs to reach high levels of broad activation in the newly grown part (Figure 2B, snapshot 2). Based on the literature, epithelia-mesenchyme interactions may play an important part in this broad activation (see Discussion), but the mechanism is largely unknown. Here, we simply assumed an extrinsic component representing the interaction with the mesenchyme. Below a certain threshold of activation, it will act to increase the concentration of the activator. Above a certain threshold, it will feedback negatively on it. This introduces an oscillatory behavior at the anterior end of the domain. Interestingly, although we do not explicitly put this in the model, the oscillatory behavior of the mesenchyme is spatially coupled to the domain growth: the transition from “no activation” to “activation” is promoted by the positive feedback from the mesenchyme, but will only happen when the domain has grown enough to escape from the influence of the inhibitor from the Turing peak (Figure 2B, snapshot 1 to snapshot 2).

In summary, our theoretical model based on *Edar* expression involves activation-inhibition mechanisms in the dental epithelium, coupled with periodic activation of the growing dental epithelium.

### A developmental palimpsest occurs for MS and R2 patterning and can be modeled with a regime of traveling wave

Next, we focused on the dynamics of *Edar* expression during the complex chain of patterning events (schematized in 1 A), that precedes the formation of the M1 signaling center, also known as the PEK (yellow in 1A/1C). The dynamics was partly similar to that observed for M2 patterning, although MS and R2 signaling centers fail to drive cap formation and to form a tooth (Prochazka et al. 2010 and figure 2A). Indeed, broad *Edar* expression restricts to these signaling centers, in late 12.5 embryos and early 13.5 embryos, respectively (corresponding to Shh-signaling center at 12.7 and 13.3 dpc in Prochazka et al. 2010 and figure 2A). Following the restriction, *Edar* expression starts again from the anterior part of the dental epithelium (white arrowhead). However, contrary to the situation in M2, for which the wave of *Edar* expression stops at a distance from the M1 area (figure 1C, 15.0 dpc), in both cases, it invaded the whole dental epithelium including its anterior most part, thus erasing the previous Turing pattern, made by MS and R2 signaling centers (figure 3A and 3B 12.5 dpc and figure 3C 13.5 pdc). In the case of MS, a new signaling center is formed following the wave: it is R2 signaling center (according to DiI tracing in Prochazka et al. 2010, it is just slightly posterior to MS). In the case of R2, two signaling centers are seen following the wave: R2 recovers and the early M1 signaling center is newly formed (although soon later, a single domain is formed see Fig 3C, 14.0 dpc, this will be discussed in detail below). We call this phenomenon a developmental palimpsest, because a palimpsest is a manuscript page that has been scraped or washed off to be used again for a novel text: here, a first Turing pattern (a first text) is erased by the *Edar* expression wave (text scraping) and a new pattern is formed (novel text).

**Figure 3.**
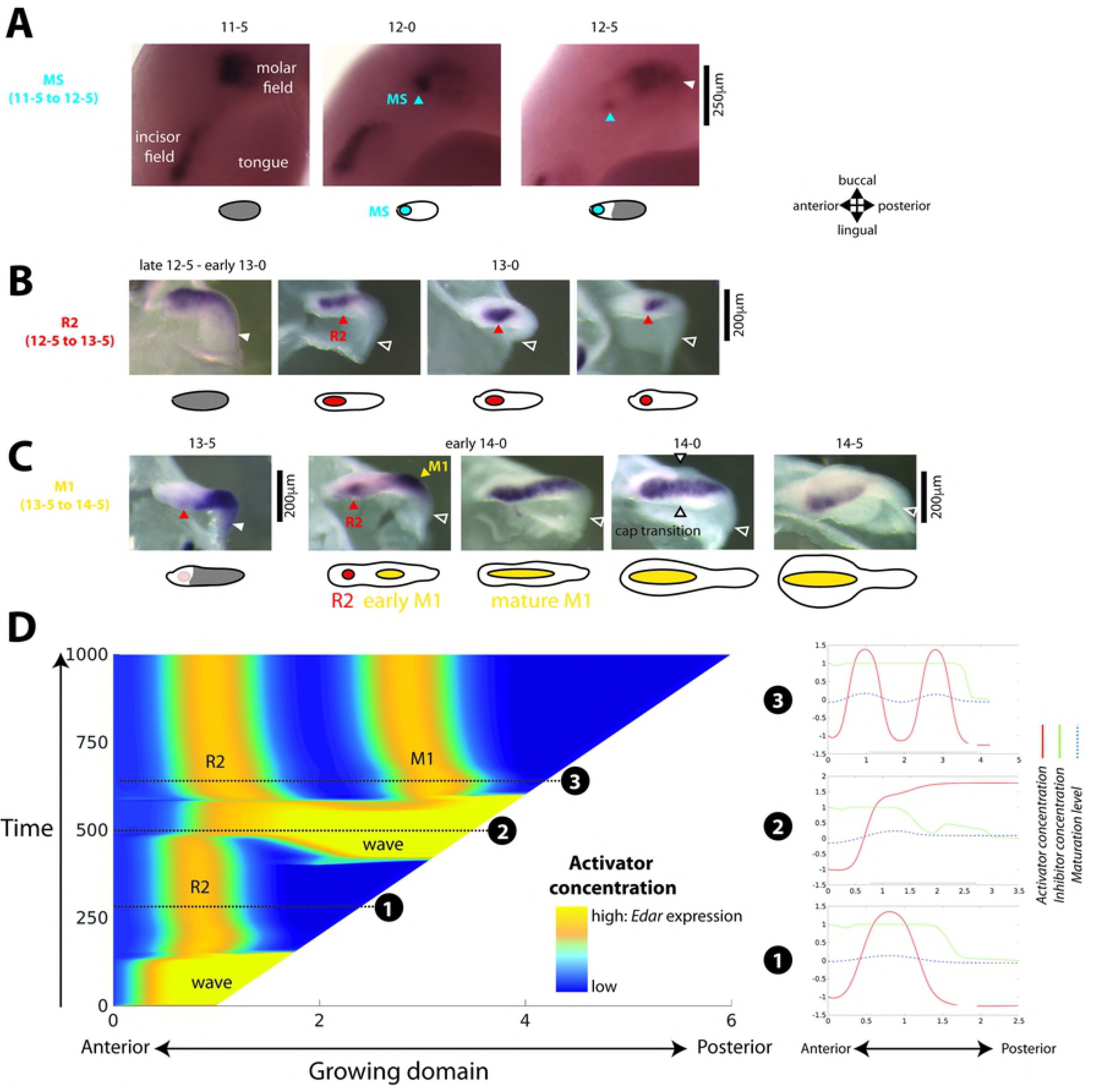
**Pattern erasing (“developmental palimpsest”) in the early dental epithelium can be modeled with a travelling wave** A, B, C - *in situ* hybridization with an *Edar* probe of whole mandibles (only one half is shown, A) or isolated dental epithelia (B, C) from 11.5 dpc to 14.5 dpc. Pictures are shown with an accompanying scheme where *Edar* broad expression is shown in grey and *Edar* focused expression is shown in color. Several rounds of *Edar* regulation are observed, culminating with the formation of a signaling center. The first pattern (MS signaling center, A) is erased by a new wave of *Edar* expression (late 12.5–13.0), resuming in a second pattern (R2 signaling center, B, D), that in turn is erased by another wave of *Edar* expression (13.5). A pattern with 2 spots is transiently observed that finally resumes in a large spot corresponding to M1 signaling center (C, D). D Pattern erasiure can be observed in a modified model, where we introduce an asymmetry in the bistable regime. The left panel shows the chronogramme of a representative simulation (concentration of activator as a function of space (x) and time (y). Snapshots taken at three different timepoints are shown (red: activator concentration, blue: inhibitor concentration, green: domain maturation). Snapshot 1 corresponds to R2 peak (13.0 dpc in panel B). Snapshot 2 shows the activation wave invading R2 domain (13.5 dpc in panel C). Snapshot 3 shows recovery of the R2 peak, together with the newly formed M1 peak (early 14.0 dpc in panel C). The movie corresponding to the snapshots is available as supplementary material 3.

In our model, we could reproduce this behavior by imposing to the immature dental epithelium a bistable regime with inactive and active stable states. This is a generic reaction-diffusion mechanism for wave propagation. A representative simulation of this modified model is shown in Figure 3D (see also the movie as supplementary material 3). The active state is more stable, so that once activation is primed in the anterior part, a wave activates progressively the inactive area (Figure 3D, snapshot 2). It is important to notice that, before wave initiation, the immature area is maintained naturally in the (less) stable inactive state under the influence of the Turing initial peak. The wave can even propagate into the mature tissue and erase the previously formed Turing pattern (Figure 3D, snapshot 1 and 2). Then, as a consequence of tissue maturation subsequent to wave activation, two activation peaks are formed by a secondary Turing patterning (Figure 3D, snapshot 3). This would correspond to R2 recovery and the newly formed M1 signaling center. For this palimpsest to occur, the wave that initiates in the immature (bistable) domain should interact with the stable pattern in the mature Turing domain. Both the activation wave and the Turing peak feature stability. As such, understanding their interaction is far from trivial. Several conditions must be fulfilled in order to observe a palimpsest in the numerical tests, which are reviewed in Supplementary Mat S3. In particular, we found that auto-inhibition, if increased in the bistable regime, strengthened the wave and favored the palimpsest. We also found that it is sensitive to the temporal dynamics. It requires a suitable synchronization between domain growth, anterior activation, wave speed, and maturation rate.

### Increasing inhibition in the model can explain the counter-intuitive rescue of R2 signalling center in *Edar^DlJ^* mutant

Most of the numerous mutants where a premolar-like tooth forms, supposedly from R2, have larger or simply normally sized molar teeth. The teeth of loss of function mutants for the Eda pathway are poorly grown, yet a premolar-like tooth can form. A rescue of R2 is counter-intuitive in such context, therefore, we decided to re-examine one of these mutants (*Edar^DownlessJ^*, abreviated *Edar^DlJ^*) in the light of *Edar* dynamics and the present model.

First, we looked at the dynamics of *Edar* expression in *Edar^DlJ^* mutant, to check if R2 was indeed rescued in this mutant. This mutant encodes for a defective Edar protein due to a single amino-acid change, but the gene is still transcribed. In contrast with the mutant epidermis, in which *Edar* regulation is lost (*Edar* expression stays at uniform low levels in the epidermis) and hair fails to form (56), we still observe *Edar* restriction to tooth signaling centers (Figure 4A), consistent with teeth being formed. However, the dynamics of activation-inhibition mechanisms was modified in this mutant. We observed high variability between embryonic tooth rows (including left/right), in line with the high phenotypic variability seen in adults with losses of function for the Eda pathway (teeth rows with 2 or 3 or in rare cases 4 teeth) (50–52). In the lower jaw (Figure 4A), we did not find obvious differences early in 12.5 dpc *Edar^DlJ-/-^* embryos as compared with Edar^D1J+/+^ embryos, all of them exhibiting restriction of *Edar* to MS signaling center that is also stained by *Shh* expression (not shown). In both cases, wild type and mutant, *Edar* expression is next found in the whole dental lamina (13.0 dpc, Figure 4A). However, no restriction was observed in *Edar^DlJ-/-^* 13.5 dpc embryos as normally seen in their wild type counterpart (FVB background). Homogenous *Edar* expression was still observed in most 14.0 dpc *Edar^DlJ-/-^* embryos at a time when homogenous *Edar* expression is again observed in *Edar^DlJ+/+^* embryos. From 14.5 dpc, a restriction to a signaling center was observed in most embryos (here named T1 PEK, with or without expression in the “tail”), while others still display more or less continuous *Edar* expression. We noticed that this signaling center in the mutant is found more posteriorly in the jaw than is the R2 signaling center (Figure 4B). At 15.0 dpc, we see either one signaling center with *Edar* expression in the tail or two signaling centers (named T1 and T2). Possibly the later case is due to approximately simultaneous patterning of two signaling centers from a dental epithelium that was showing continuous expression in the previous stage. At 15.5, T1 has developed into tooth germ of different sizes, from a very small one to a tooth germ equivalent in size to T2. To conclude, our results show that R2 patterning is both postponed and displaced posteriorly in the *Edar^DlJ-/-^* mutant and the resulting signaling center persists to form a tooth germ of variable size. Why then is R2 rescued in the context of poorly grown teeth?

**Figure 4.**
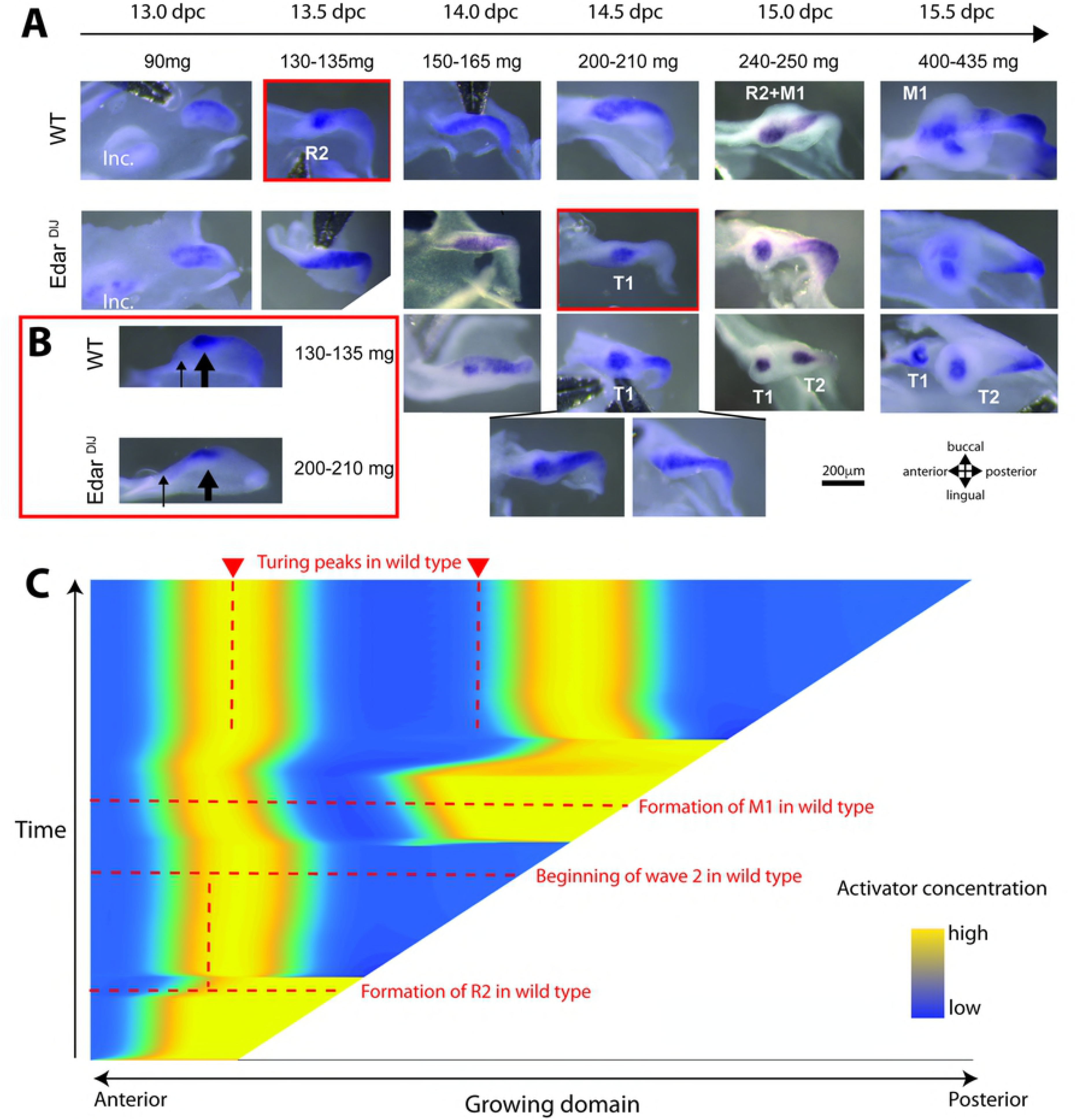
**the changes in signaling center patterning observed in the *Edar^DlJ^* mutant are predicted by the model** A- In situ hybridization with an *Edar* probe in dissociated lower jaw epitheliae from *Edat^DlJ/DlJ^* mutant and its wildtype background (top views), for embryos of similar weight class between 13.0 and 15.5 dpc. Patterning of the first signaling center (MS) is correct (not shown), but patterning of the second signaling center (R2, here called T1 in the mutant) is delayed with variability (embryos >200mg). This T1 signaling center is not erased and persists. Inc: incisors. B- A side view of pictures outlined in red in A, showing that T1 signaling center is found more anteriorly in the invaginated dental epithelium of the mutant as compared to the wild type. C- Same simulation as in figure 3, but with increased inhibition. The position and timing of Turing peak formation in wild type (as in figure 3E) are indicated with red dashlines. Increasing the inhibition (lowering auto-inhibition) is sufficient to abolish the palimpsest and stabilize the first and the second peak.

We next used our model to explain this non-intuitive observation. Eda pathway loss of function has been shown to increase inhibition in other appendages (Mou et al. 2006; Harjunmaa et al. 2014), which is consistent here with T1 signaling center being patterned further away from the anterior end of the dental epithelium. Therefore, we analyzed numerically the effect of stronger inhibition (by decreasing auto-inhibition rate). Interestingly, this was sufficient to recover qualitative behaviors consistent with the data: 1-T1 and T2 are formed later than R2 and M1 (red dashed line). 2-T1 is displaced anteriorly (blue dashed line) 3 - the wave no longer destabilizes T1: a Turing pattern is formed that is not subjected to a palimpsest. These results suggest two things. First, that a new wave of activation associated with the formation of the next signaling center will naturally destabilize any pre-existing signaling center, if inhibition from this pre-existing center is weak enough. Second, because we do not impose any difference between signaling centers, this shows that R2 is not intrinsically defective, but actively outcompeted by the activation wave associated with M1 formation. This is a major output of our modeling effort but it is in contradiction with the previous hypothesis, in which the R2 signaling center is considered to be intrinsically defective. Therefore, we decided to directly test this hypothesis.

### The anterior part of the dental lamina (R2) is capable of forming a tooth - if separated from the anterior part

If the anterior part of the dental epithelium is intrinsically defective for tooth formation, it should not be able to give rise to a fully develop tooth when removed from the early M1 signaling center. On the contrary, if the anterior part is not intrinsically defective, but normally outcompeted by the M1 as we suggest, a tooth should be able to develop from it when removed from M1 influence. To test this, we cut the anterior part (R2 part) from the anterior part (early M1 signaling center and anterior tail). As expected from our predictions, the anterior part developed into a fully growing tooth. Remarkably, the timing of development is advanced compared to the anterior part by 1 day, in accordance with the R2 signaling center having been patterned earlier than the M1 one (Figure 5).

**Figure 5.**
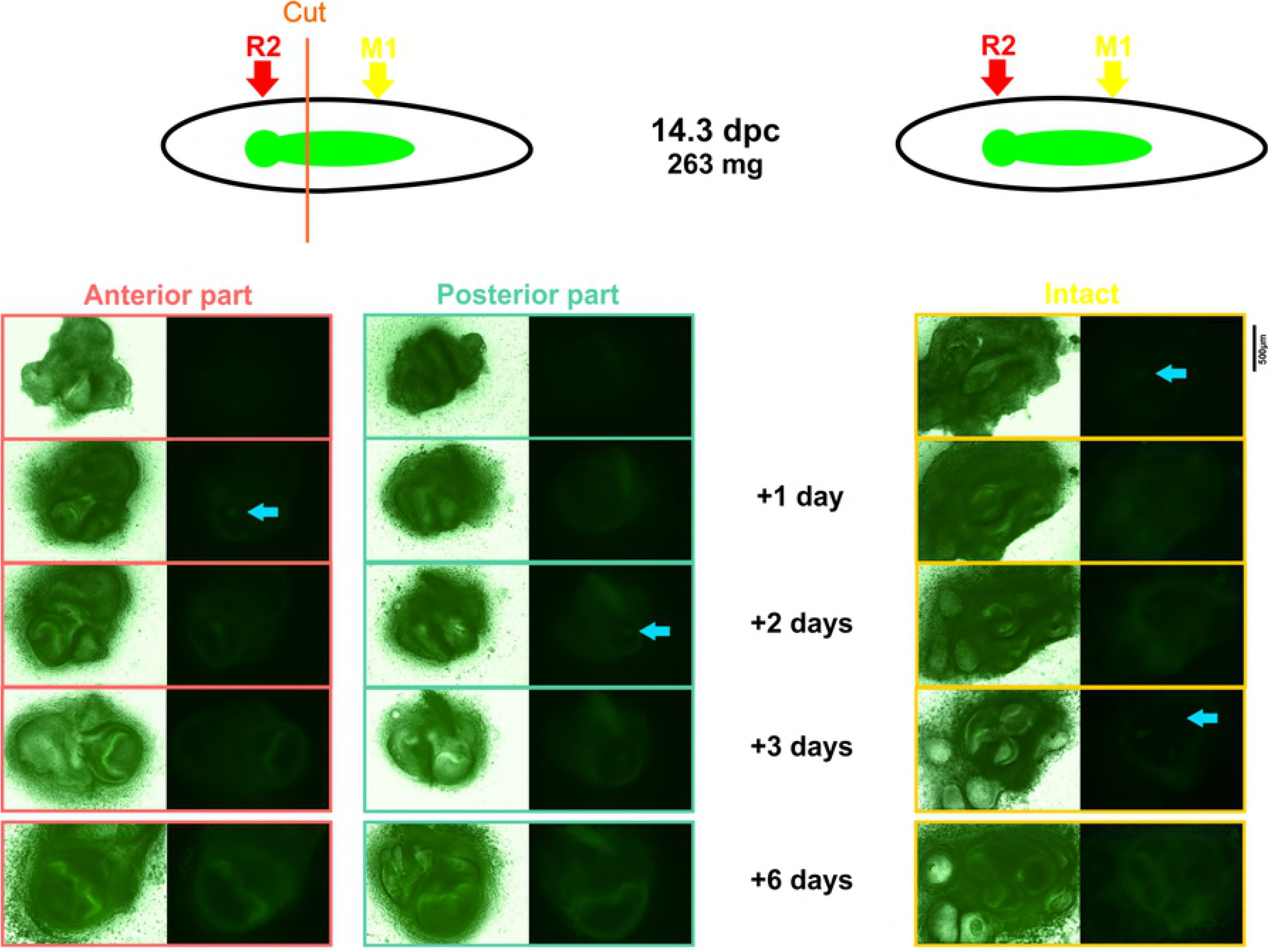
**The anterior part of the molar field corresponding to R2 signaling center can develop to a tooth germ when separated from the anterior part** Left: The developing molar region was dissected from 14.3 dpc embryos and the anterior most part corresponding to R2 signaling center was separated from the rest, including the early M1 signaling center and the anterior “tail”. The following day, a signaling center was recovered in the anterior part and formed a tooth germ (6 days cultivation). After two days, a signaling center formed in the anterior part and formed a similarly sized tooth as the anterior part. In the control experiment (right), the second molar does not form before the third day.

Taken together, these results are consistent with a model in which the R2 region is fully competent for tooth formation, but is actively outcompeted by the forming M1, resulting in the developmental palimpsest effect described.

### The formation of a large signaling center depends on Edar activity

We then focused on another feature of the mouse dental row: the incorporation of R2 into M1 which is hypothesized to play a crucial role for the formation of the anterior part of M1, both during development and evolution (30,33). This corresponds to another curious behavior of *Edar* expression dynamics: the fusion of R2 and early M1 signaling centers soon after recovering from the palimpsest (Figure 3C). We examined the 13.5–14.5 dpc period, which corresponds to M1 PEK formation, in detail. To do so we followed in parallel the dynamics of *Edar* expression, *Shh* expression (a recognized marker of tooth signaling centers) and Wnt pathway activity (monitored by the TOPGAL reporter) (Figure 6A). In late 13.5 dpc/ early 14.0 dpc embryos, *Shh* expression and TOPGAL X-gal staining reveals that the M1 signaling center starts to form. Some faint *Shh* expression is occasionally seen in R2, while X-gal staining persists there, presumably in part due to B-galactosidase long half-life. At this stage, *Edar* is also focused in R2 and M1 signaling centers, yet low expression can also be seen around. In slightly older embryos, robust *Edar* expression is seen in a domain spanning the two signaling centers, and aligned with the barely formed cervical loops. This *Edar* expression is followed by anterior expansion of *Shh* expression, that finally spans the position of former R2 and early M1 signaling centers (as shown in Prochazka et al. 2010). We also observed upregulation of TOPGAL activity during the same period. All together, these results show that R2 and early M1 signaling centers are re-patterned as a single large signaling center highlighted by Edar expression. This early event prefigures TOPGAL activity and *Shh* expression relocalization in a large signaling center.

**Figure 6.**
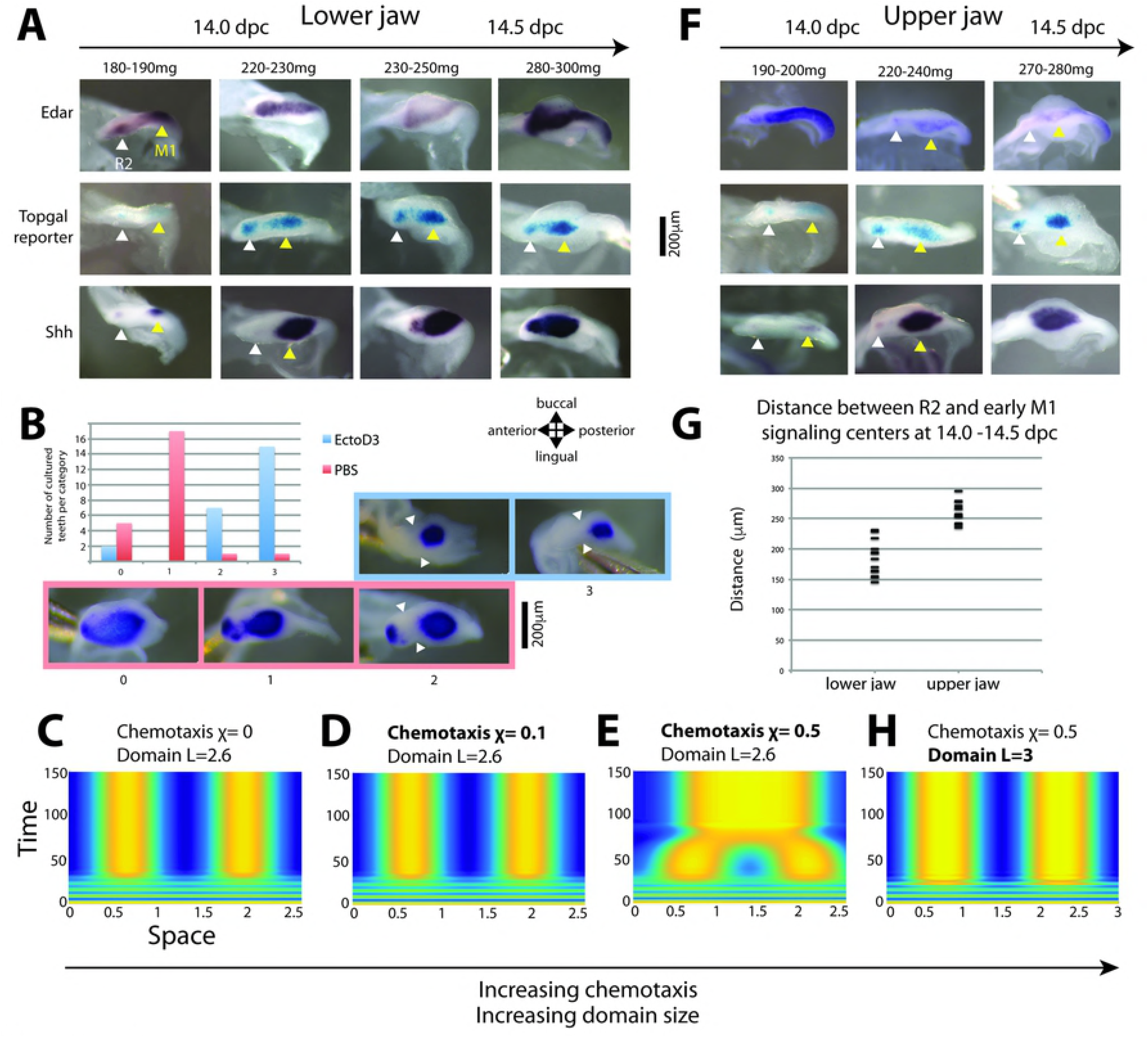
**The formation of a large fused PEK depends on Edar signaling, possibly through chemotaxis A,F:-** Dynamics of *Edar* (upper) and *Shh* (lower) expression and TOPGAL reporter X-gal staining (middle) during the 14.0 to 14.5 dpc period when M1 PEK forms, in lower (A) and upper (F) jaw of CD1 mice. In both jaws, *Edar* expression spans both R2 and M1 region before becoming restricted. In upper jaw, and very transiently in the lower jaw, expression restricts again to R2 plus the newly formed M1 signaling center (220–240 mg and 180–190 mg, respectively). In lower jaw, expression restricts to a large domain including both R2 and early M1 signaling center (from 220–230 mg). *In situ* hybridization with *Edar* or *Shh* probes and X-gal stainings were performed on dissociated lower or upper jaw epitheliae. The embryonic weights used to stage embryos are shown. Staining concentrated in R2 signaling centers is pointed with a white arrowhead, while M1 signaling centers are pointed with a yellow arrowhead. B -Lower molar rows were put into culture at 13.0 dpc and following 2 hours of recovery, they were treated with an Eda blocking antibody (EctoD3) or with mock for 40h. The dissection process tends to interfere with the formation of a large signaling center: in most samples, low *Shh* expression is seen between R2 and early M1 signaling center; however, R2 expression is always maintained. In EctoD3-treated samples, R2 expression is lost and only a small posterior M1 signaling center is found. C, D, E, H -Numerical simulation of a simple Turing model augmented with chemotaxis in a fixed, matured domain with a simple configuration consisting in two Turing peaks C: in absence of chemotaxis (χ=0), two Turing peaks form in a domain of length L=2.6. D, E: in a model where the activator stimulates the production of a chemoattractant (over a certain threshold), Turing peaks form first, and later chemotaxis fuse the two peaks in a single large peak (χ=0.5, E). However, the fusion requires sufficient activity of the chemoattractant (%=0.1, D). H On a larger domain (L=3), a Turing pattern with a larger wavelength is chosen, similar levels of chemotaxis as in D (χ=0.5) are no longer sufficient to fuse the peaks. G - Distances between R2 and M1 signaling center as measured on TOPGAL dissociated epithelia at 14.0 dpc in lower and upper jaw. R2 and M1 signaling centers form closer in lower jaw than in upper jaw.

Because *Edar* expression prefigures the anterior expansion of *Shh* expression domain and *Edar* has been shown to regulate *Shh* (42,57,58), we wanted to test if Edar signaling is necessary for anterior expansion and the formation of a large M1 PEK. To specifically test this, we dissected 13.0 dpc lower molar regions, when R2 has already formed, and cultured them for 48h with or without an interfering antibody, so that we knock-down Edar signaling in the next period of M1 PEK formation. We then visualized *Shh* expression on isolated epithelia. In untreated samples, we occasionally observed a large M1 PEK similar to *in vivo* samples (Figure 6B, state 0), but most often it was split in two spots (corresponding to R2 and an extended early M1 expression) bridged by a narrow domain of *Shh* expression (Figure 6B, state 1). This can be explained by the fact that the dissection process could change the activation-inhibition balance in favor of inhibition (as proposed in (59)), which according to the predictions of our model, should favor R2 persistence. In treated samples, we mostly recover a small, very posterior signaling center, which corresponds to early M1 *Shh* expression (Figure 6B, state 2 and 3). Moreover, the dental epithelium is morphologically different, showing a bud, followed by a small cap and a “tail”. Thus, in the absence of Edar signaling, *Shh* expression is lost in the R2 region and only a small PEK forms, which is equivalent in size and position to the early M1 signaling center, and which drives cap transition there. Taken together, our results show that Edar signaling is essential for the formation of a large PEK that encompass R2 and early M1 signaling centers.

### Coupling chemotaxis to the Turing system reproduces biological variability in signaling center fusion

Recent studies have pointed out that chemotaxis may plays a role in the formation of tooth and hair placodes (10,45,46) and the Eda pathway activated centripetal migration in the placodal epithelium (45). We noticed that the TOPGAL stainings tend to contract in the antero-posterior direction as the M1 signaling center matures and the distance between R2 and early M1 signaling center decreases (compare 14.0 and 14.5 dpc samples in figure 6A). This suggested us that cell movements may take part in the formation of the large signaling center.

To evaluate this possibility, we incorporated cell motion through chemotaxis in a simple Turing system producing two peaks, thus starting with the situation when R2 and M1 signaling center co-exist (Figure 6C). We assumed that the chemoattractant pattern corresponds to the activator pattern (*e.g*. cells move towards regions of higher activator concentration). Cell aggregation mediated by chemotaxis requires a positive feedback loop between cell density and chemoattractant concentration (18,19). In our setting, without addition of extra molecular entities, this feedback loop can act either directly on the activator-chemoattractant (as in the classical Keller-Segel system, (19)), or *via* down-regulation of the inhibitor (*e.g*. higher cell density affects negatively inhibitor concentration). The latter configuration produced the expected behavior: signaling centers form and secondarily they fuse in a single large signaling center (Figure 6E). This is consistent with the intuitive idea that fusion requires sufficiently long range communication between the two Turing peaks, and thus a feedback on the long-range diffusing species (in our case the inhibitor). Reducing chemotaxis efficiency resulted in the lack of fusion, consistent with our experiments reducing Edar signaling, and presumably, chemotaxis (Figure 6D).

Nonetheless, adding chemotaxis to our system does not necessarily result in the fusion into a single spot. In our *in silico* simulations, we observed that the transition between fusion and the absence of fusion depends on various parameters. The reason is that the activator-chemoattractant has a direct positive effect on the inhibitor, but an indirect negative effect on the same inhibitor by means of cell recruitment. These ambivalent effects make chemotaxis able to compensate the segregation due to Turing patterning in some situations, or reinforce it in other situations. For example, we observed that chemotaxis can favor pattern formation where the Turing system fails to produce a pattern alone. This suggests us that chemotaxis may be part of the normal formation of tooth signaling centers, even when they stay separated.

In line with this, we noticed that there is no fusion in the upper jaw (Figure 6F), where the distance between signaling centers is initially larger by about 30% (Figure 6F and G). A small increase of the domain size, as seen between lower and upper jaw, increased the Turing wavelength and was sufficient to abolish the fusion under the same chemotaxis efficiency (Figure 6H, a 15% increase is sufficient with our parameters setting).

In conclusion, the non-trivial interaction between chemotaxis and a Turing system appears to be a plausible mechanism to explain the variability in the dynamics of tooth signaling centers in lower jaw, upper jaw and various mutants, including *Edar* loss of function.

## Discussion

In this study, we have revealed the highly complex and dynamic behavior of signaling centers responsible for tooth patterning in the mouse jaw. Patterning is usually seen as a directional process, rather than a dynamic process that could take circuitous routes. However, we show that patterning of the first molar involves what we called a developmental palimpsest, where patterns are established, erased or remodeled to give rise to new patterns. Using a mathematical approach, we show that these behaviors (pattern erasing, recovery, rescue, fusion) despite seeming to be complex can be be produced by the activity of simple mechanisms (a Turing pair with two regimes, as well as chemotaxis).

### From similarities in *Edar* expression dynamics to differences between hair and tooth patterning

In this study, we have revealed the highly dynamic expression of *Edar* in the developing molar row. This dynamic is superficially similar to that seen during hair patterning. This is not surprising since hair and tooth patterning share many common features (Mikkola, 2007; Biggs and Mikkola, 2014), making their comparison highly instructive. Below, we compare these two systems in light of our results.

In teeth, as in hair, *Edar* expression becomes restricted to the signaling center as it is patterned. We have noticed however two substantial differences: i) In skin, the initial basal levels of *Edar* are upregulated in the placode and downregulated in its vicinity. This is thought to be pivotal for placode patterning, where Eda signaling is necessary to stabilize and refine an otherwise labile Wnt-dependent placode prepattern (11,43). In the molar field, *Edar* expression in the dental lamina reaches levels pretty similar to restricted expression in the signaling center, suggesting that *Edar* is actively stimulated in both cases. This upregulation may rely on ActivinβA (stimulating *Edar* expression in tooth cultures, (Laurikkala et al. 2001) and the Wnt pathway (which plays a central role in tooth formation, and is involved in *Edar* basal expression in hair (11,43)). Downregulation may rely on the BMP pathway, as in hair ((11,43)). ii) We show that this regulation still occurs in the *Edar* mutant, a major difference with hair, for which the regulation does not occur and uniform basal levels of *Edar* expression are maintained in the absence of Eda signaling (11). Self-activation of the pathway thus plays a more minor role, if any, in teeth.

We believe that these differences on *Edar* regulation may reflect differences in the balance of the different processes participating in hair and tooth formation. For the formation of hair placodes, Turing-like mechanisms establish a noisy pre-pattern, with local sources of FGF signaling. Mesenchyme condensation towards these sources then refines and reinforces the pattern (Glover 2017). The mesenchyme is also able of autonomous self-organisation, but this is masked by the pre-pattern imposed by the epithelium.

The formation of tooth signaling centers seems to rely on a different equilibrium between the two tissues. The formation of a PEK is highly dependent on mesenchyme condensation, as seen in bud-arrested tooth germs where condensation fails (29). Modeling the gene network of epithelium-mesenchyme interactions in teeth also lead to the suggestion that the two tissues work in concert, rather than one dominating the other (O′Connell *et al*, 2012). These intrinsic differences may explain why *Edar* loss of function abolishes pattern formation in the epithelium-dominated context of hair formation, but only results in spatio-temporal modifications, in the more balanced context of tooth formation.

### A model for sequential patterning of signaling centers in the dental epithelium

In this study, we assume that the complex spatio-temporal changes in *Edar* expression highlight waves of activation in the dental epithelium. Each of these waves resumes with the patterning of a signaling center and they are reiterated upon posterior growth of the dental lamina. We note that this growth zone could be the *Sox2* positive region shown in (23). We built a reaction-diffusion mathematical model to describe this behavior. In this macroscopic model, molecules are treated as a continuum, and set on a 1-dimensional space to model the antero-posterior dimension of molar row formation. We also chose to consider only 2 types of Turing in-phase molecules, corresponding to an activator and an inhibitor. This is of course a high level of abstraction. Tooth genetics has revealed many molecules from the epithelium or the mesenchyme that could participate in the activation-inhibition mechanisms (with both in-phase and out-of-phase patterns), but it was not our purpose here to identify these molecules. Instead, our modeling effort aimed at providing a theoretical framework for sequential tooth formation. Moreover, although our model explicitly aims to describe activation in the epithelium (*Edar* dynamics), this does not mean that the activator-inhibitor couple in our model should be seen as an abstraction for Turing reactions in the epithelium only. We do not rule out that our model could synthesize the activation-inhibition reactions arising from epithelia-mesenchymal interactions and giving rise to the Turing pattern.

We focused on qualitative insights and assessed robustness of pattern formation and developmental palimpsest in our model. We found a suitable model parametrization, and tested its sensitivity with respect to patterning (see Supp Mat for details). Although the results are generally robust enough to moderate parameter changes (10%-50%), it is interesting that the developmental palimpsest can be abolished in many ways, changing auto-inhibition but also temporal dynamics and synchronicity between events. This is consistent with the marked tendency of molar row development towards supplementary molar formation: it can happen in mutants from many different pathways, and moreover it often occurs without major changes in other aspects of tooth development.

Our model explicitly assumes that the mesenchyme is responsible for periodic activation priming the newly grown epithelium. This dependence is consistent with a body of evidence showing that mesenchyme activity is necessary for the induction of primary enamel knot formation and sequential tooth formation (61,28,27). We also know that mesenchyme activity depends on the msx1-Bmp4 feedback loop (62–64), which is itself dependent on a mechanical signal provided by mesenchyme condensation (Mammoto et al.)(29). When this loop is defective, sequential tooth formation can stop at different stages from no tooth forming, only one, or only two instead of three (65,63). It can also simply stall until adequate levels of Bmp4 signaling are reached, as seen in the barx1 mutant (66). The mesenchymal Bmp4 signal is part of a Wnt-Bmp regulatory network whose integration drives signaling center formation (28). It is also known that the mesenchyme produces ActivinβA, a potent inducer of both tooth formation (Kavanagh et al. 2007) and *Edar* expression. In the absence of further knowledge about how the mesenchyme could prime the waves of activation observed in the epithelium, we introduced in our model an extrinsic component representing the interaction with the mesenchyme, and chose a parsimonious way to provide it an oscillatory behavior. For this, we assumed that the mesenchyme activity is stimulated by the activator and feedbacks on it in the newly grown area. Below a certain threshold, it will act to increase the concentration of the activator. Above a certain threshold, it will act to decrease the concentration of the activator.

Another interesting feature of our model is that inhibition from the Turing spot locks the bistable system of the newly grown epithelium in the “no activation” state. This means that, in the absence of a wave of activation triggered by the mesenchyme, sequential addition will stop, in contrast with a standard Turing system in a growing field. This is consistent with mutants in the bmp4-msx1 axis where sequential addition resumes after M1 or M2 formation. Experimental approaches will be needed to determine the mechanisms enabling periodicity in our system, and to implement this in the model. Further investigation within our modeling framework should involve two separated compartments for epithelium and mesenchyme. It is expected that regulation feedbacks and delayed growth can trigger intrinsic oscillations if the mesenchyme can escape the “locked” inactivated state under long-range inhibition.

Our model shares some similarities with models of somitogenesis. First of all, almost all somitogenesis models include a clock driving gene expression oscillations, forming traveling waves moving through the tissue (e.g. (14)). Even cells isolated from the presomitic mesoderm exhibit oscillations (67). However, whether such a *bona fide* molecular oscillator will be found in the tooth system remains an open question.

We note that tissue-scale oscillations have been observed in limbs, whose development share similarities with that of epithelial appendages including teeth (68). We also envision other possibilities relying on tissue properties rather than cell properties, for example emerging from the cross-talk between the epithelium and the mesenchyme (as suggested above).

Second, in the long-prevailing models of somitogenesis, the clock is combined with a gradient of Fgf/Wnt signaling, that maintains the oscillations in the posterior part and determines the position where the traveling wave is frozen into a stationary pattern, which will define somite boundaries (69). Our model does not comprise such positional information. In a more recent model of somitogenesis, traveling wave and pattern formation are produced by a Turing pair with a non-diffusing activator and a diffusing inhibitor (Cotterel et al. 2015). Pattern formation arises when the traveling wave breaks next to the previously formed stripe (that acts as a stable source of inhibitor), and local interactions in this region promote activator increase to form a new Turing stripe in the vicinity of the previous stripe. This model shares an obvious similarity with our model: a Turing pair exhibits different behaviors (oscillatory with traveling wave / Turing in the Cotterel model versus bistable with traveling wave/Turing in our model) along the antero-posterior axis. However, the switch between the two behaviors arises as a local emergent property next to previously formed stripes in the Cotterel model, whereas it is explicitly introduced in our model as a result of maturation. Moreover, in our model the oscillations are provided as an independent term, materializing mesenchyme function. We acknowledge that the Cotterel model might apply to the tooth system, and it will be interesting to test if the palimpsest can be obtained with such a model.

Our study also shares superficial similarities with another system showing sequential patterning: feather patterning. In this system, a priming wave of activation is observed in the epithelium, giving rise to a stripe in the chick embryo’s back, which is then broken into a spot pattern giving rise to individual feathers. Pattern formation, in the model by Painter et al., relies on chemotaxy rather than reaction-diffusion (21): moderate cell aggregation drives stripe formation in the primed epithelium through a FGF-dependent positive feedback, and strong local aggregation introduces a BMP-dependent negative feedback that contributes to break the stripe into spots. The behaviors of the two systems are similar: the broad Edar expression could be compared to the priming wave/first stripe, and the formation of the signaling centers to the breaking of the stripe into spots. These models also converge conceptually. Stripe formation in the feather model, and Edar activation wave in the tooth model, mainly rely on positive feedback. Spot formation and signaling center formation both rely on the introduction of a sharper negative feedback. We take this as an indication that this sequence of activation might be a general property of epithelial appendages (feathers, hair, teeth), that can be captured by very different, non-exhaustive models. We also want to stress that our model is meant to recapitulate activation/long-range inhibition mechanisms, rather than specifically reaction-diffusion mechanisms, and we do not exclude that the biological mechanisms it captures are based on chemotaxy, as in the Painter model.

### Making and erasing patterns: a developmental palimpsest characterizes first molar formation

Lastly, we would like to emphasize how the current model successfully recapitulates a number of counter-intuitive behaviors of the system and inform us on the possible underlying mechanisms.

Previous studies had already revealed several complex behaviors in the growing dental epithelium: i) the transient patterns of MS and R2 signaling centers, supposedly vestiges of premolar signaling centers; ii) the rescue of an abortive tooth germ R2, in a large number of genetic conditions; iii) the transient co-existence of R2 and early M1 signaling center followed by their fusion in a large signaling center in the lower jaw.

The present data and our simple model suggest that these complex behaviors are the fruit of rather simple but highly dynamic interactions in the growing tooth field.

As viewed from Edar expression, the pattern constituted by MS, and later R2 signaling centers is erased to give raise to a second wave of patterning, materialized by a broad Edar expression in the dental epithelium at respectively 12.5–13.0 dpc and 13.5–14.0 dpc. This was recapitulated in the model by enabling the bistable domain to form a traveling wave, that can destabilizes a previously formed signaling center, if inhibition in the later is not too strong. Aside from recapitulating Edar expression, the travelling wave has more profound implications. Indeed, it implies a first paradigm shift that vestigial buds are not committed to abort as usually thought (for example due to their proximity with the diastema thought to serve as a source of inhibitors (70,71), or through the expression of specific molecules (72)), but rather (or on top of that) that they are actively competed by the next round of activation as the dental epithelium grows. Such a balance explains why the anterior part of the molar row is very sensitive to genetic perturbations (with many genetic conditions exhibiting a supplementary tooth there) and environmental perturbations (i.e. tooth culture in this study). It also explains why even conditions that produce a more inhibitory context than the wild type can produce such supplementary tooth. Indeed, our model predicts that if inhibition is increased (e.g. auto-inhibition is decreased), like it is commonly assumed in the Eda pathway mutants, then the Turing pattern remains (with a slightly longer wavelength), but the traveling wave is almost immediately suppressed. This is exactly what we document in Edar^DlJ^ mutants for the R2 signaling center: it forms more posteriorly, and we see no traveling wave that would erase it. Rather, it persists to form a tooth bud. Our cultivation of anterior parts of the molar field, corresponding to R2 signaling center, also show that it has the potential to fully form a tooth, but is actively competed by the M1 signaling center in the wild type situation. Consistent with our results, Li *et al*. reported that FGF8 application could rescue tooth germ development in the mouse diastema only when it was separated from the molar and incisor buds (73). In conclusion, our results extend the prevailing model (that of Kavanagh et al.), where inhibition between forming teeth is unidirectional (from M1 to M2, to M3), by showing that inhibition can be bidirectional and subtly dependent on the temporal dynamics of the system.

In the wild type, following broad *Edar* activation at 13.5–14.0, a new pattern of *Edar* restriction forms that is markedly different between the lower and upper jaw. In the lower jaw, independent R2 signaling center and early M1 signaling center are transiently seen (very transiently with *Edar* expression, for a longer time with *Shh* expression or TOPGAL activity) but they are rapidly included in a single elongated signaling center. In the upper jaw, *Edar* restricts to R2 signaling center and M1 signaling center and remains as such. Therefore, the palimpsest is observed in both jaws, enabling the co-occurrence of two signaling centers, but only in the lower jaw does some additional mechanism enable their fusion into a large signaling center. Here, we introduced chemotaxy to the model, because chemotaxy has been evidenced in hair placode formation, both at the level of the epithelium (45,46) and the mesenchyme (10,21). This was sufficient to recapitulate a number of interesting features: 1) chemotaxy changes the reaction-diffusion so that the system first makes two peaks that later fuse into a single, larger peak: this is reminiscent of the large M1 signaling center 2) this behavior is sensitive to the distance between the initial peaks, and a 15% increase was sufficient to impede fusion. The measured 30% difference between the R2-M1 distance in the lower jaw, where fusion occurs, and the R2-M1 distance in the upper jaw, where fusion does not occur, may thus be sufficient to explain difference in fate in the two jaws. 3) finally, reducing chemotaxy was sufficient to impede fusion. This may explain why in our culture system, inhibition of Edar activity impedes fusion although the distance between R2 and M1 does not seem drastically changed.

Interestingly, we show that chemotaxy plays an ambivalent role in our model. Depending on the conditions, it acts in favor or against the Turing pattern, or it is relatively neutral. We propose that this ambivalent role contributes to explain the versatility of this biological system in regard to genetical and environmental (e.g. culture) perturbations.

## Conclusion

An important lesson from the tooth system is that patterning events may be less straightforward than usually thought, and patterns may be dynamically drawn and erased or refined during embryogenesis. In other words, developmental palimpsests may be a common feature. One reason for this is historical: systems are the product of evolution, and as pointed out by F. Jacob, evolution proceeds as a tinkerer, not as an engineer (74): extant patterning mechanisms are modified versions of ancestral mechanisms, and not purposely designed from scratch. Our study is consistent with recent studies on the fine-scale temporal dynamics of gap gene patterns in dipterans (e.g. a progressive anterior shift of the gap genes pattern). As shown here for the *Edar* mutant, incorporating this dynamics into models provided a better explanation for mutant phenotypes (8,75,76). Moreover, this curious dynamics is also likely the vestige of an ancestral mode of segmentation (75,77). In summary, we believe these two systems illustrate that temporal dynamics of developmental systems needs to be studied, and moreover to be studied in the light of evolution, to fully explain how the system reacts to perturbations. Indeed, embryonic patterns can be highly dynamic and thus dynamics can be essential to the outcome of the patterning process.

## Materials and Methods

### Mouse breeding and embryo harvesting and staging

All animal experimentations were conducted under animal care procedures in accordance with the guidelines set by the European Community Council Directives. Experimental procedures were approved by an official ethical committee (CECCAPP, Lyon; # ENS-2009-027 et # ENS-2012-046).

The CD1 mice were purchased from the Charles River (Germany). Other mice have been bred at the PBES (Lyon). TOPGAL mice (Tg(Fos-lacZ)34Efu) carrying three LEF1/TCF1 binding sites fused to a minimal *c-fos* promoter driving *lacZ* expression were backcrossed against CD1 mice for 10 generations {DasGupta, 1999 #6}. TOPGAL positive mice were screened by standard lacZ staining performed on the first phalange cut from PN4-PN7 newborns. The *Edar^dl-J^* mice (FVB background) were obtained from Paul Overbeek. They carry a G to A transition mutation causing a glutamate to lysine substitution in the death domain of the Edar protein (E379K, Headon and Overbeek, 1999). The strain was maintained by crossing heterozygotes with homozygotes, and wild type and *Edar^dl-J/dl-J^* mice used in experiments were derived from this same stock. The Eda^Tabby^ mice carry a X-linked null mutation in the *Eda* locus with a deletion of the first exon (Probst 2008). The colony was established by inbreeding from a mating pair (B6CBACa Aw-J/A-Eda^Ta^/J-XO female and B6CBACa Aw-J/A male) obtained by the Jackson Laboratory (Bar Harbor, Maine).

In order to harvest embryos every 12h hours of development, mice were kept under two different day-night light cycles. Mice were mated overnight and vaginal plugs were detected in the next morning, noon being indicated as the embryonic day (ED) 0.5. Pregnant mice were killed by cervical dislocation and embryos were harvested and weighted as described earlier {Peterka, 2002 #30; Prochazka et al 2010}.

For tooth culture experiments, CD1 females were crossed with males carrying the fusion protein Shh-EGFP (Enhanced Green Fluorescent Protein) and Cre recombinase from the endogenous *Shh* locus (B6.Cg-*Shh^tmi(EGFP/cre)Cjt^/J)* what enabled determination of *Shh* expression using fluorescence. The breeding pairs *B6.Cg-Shh^tmi(EGFP/cre)Cjt^/J* were purchased from the Jackson Laboratory (Maine, USA). The mice were genotyped using the Jackson Laboratory’s protocols. All used animals were fed and watered ad libitum. Housing of animals and experiments were carried out in strict accordance with the national and international guidelines (ID 39/2009) and under supervision of the Professional committee for guarantee of good life-conditions of experimental animals at the Institute of Experimental Medicine, the Czech Academy of Sciences, Prague, Czech Republic and approved by the Expert Committee at the Academy of Sciences of the Czech Republic (permit number: 81/2017).

#### Mandible epithelium dissociations

Mandibles and maxilla were dissected in Hank’s medium and treated with Dispase II (Roche) 10mg/ml at 37°C for 1 to 2h20 depending on embryonic stage. Epithelium was carefully pealed and fixed in PFA 4%.

#### Whole mount In situ hybridization (WISH) and X-gal staining

Embryonic mandibles, maxilla or dissociated epithelia were fixed in 4% PFA solution over night at 4°C and *In situ* hybridization was done according to a standard protocol. DIG RNA probes were transcribed *in vitro* from plasmids described elsewhere: *Shh* {Echelard, 1993 #1}, *Edar* {Laurikkala, 2001 #75}. TOPGAL embryonic mandibles or dissociated epithelia were fixed in 4% PFA for 15 minutes only and stained with X-gal according to a standard protocol. The samples were documented on a Zeiss LUMAR stereomicroscope with a CCD CoolSNAP camera (PLATIM, IFR128, Lyon) or on a LEICA MFA205 stereomicroscope with a DFC450 camera (IGFL, Lyon).

#### Organotypic culture and treatments

The lower molar region of 13.0 embryos were dissected and cultured according to methods described in Kavanagh et al. 2009. Following a period of 2 hours of recovery, the medium was changed for a new medium supplemented with 5ug/ml Eda interfering antibody (ectoD3; (78)). Tooth culture was stopped at 40h and epithelium were dissociated for 15–30 minutes Dispase II (Roche) 10mg/ml at 37°C.

#### In vitro cultures of anterior and posterior parts of M1 tooth primordium

M1 tooth germs of *ShhEGFP^+^* mouse embryos at 14.3 dpc were dissected from embryonic lower jaw and cut to anterior and posterior part. Both parts were cultured separately on PET track-etched membrane. Contralateral intact M1 dissected tooth germs from the same specimen were used as control. Cultures were photographed using inverted fluorescent microscope Leica AF6000 (Leica Microsystems GmbH, Germany) daily from day of dissection to day 6 of culture.

### Mathematical modeling

The model is described in supplementary material 1.

## Acknowledgements

We thank Denis Headon for providing Edar^DlJ^ mutant, Irma Thesleff for providing *Edar* probe. We acknowledge the contribution of SFR Biosciences (UMS3444/CNRS, US8/Inserm, ENS de Lyon, UCBL) facilities: PLATIM and PBES and warmly thank their respective staff for their constant help. We thank Nicolas Gadot and the ANIPATH facility for related work. We thank Neal Anthwal for helpful comments on the manuscript.

## Supporting information

**Supplementary material 1 (.pdf):** a detailed description of the model together with simulations under different parameter ranges.

**Supplementary material 2:** a movie (.gif) of the simulation shown in figure 2. The evolution of activator concentration (red), inhibitor concentration (blue) and domain maturation (green) are shown as the domain grows.

**Supplementary material 3:** a movie (.gif) of the simulation shown in figure 3. The evolution of activator concentration (red), inhibitor concentration (blue) and domain maturation (green) are shown as the domain grows.

**Supplementary material 4:** a movie (.gif) of the simulation shown in figure 4. The evolution of activator concentration (red), inhibitor concentration (blue) and domain maturation (green) are shown as the domain grows.

